# IL8 Drives an Elaborate Signal Transduction Process for CD38 to Produce NAADP from NAAD and NADP^+^ in Endolysosomes to Effect Cell Migration

**DOI:** 10.1101/2020.03.24.004739

**Authors:** Tae-Sik Nam, Dae-Ryoung Park, So-Young Rah, Tae-Gyu Woo, Hun Taeg Chung, Charles Brenner, Uh-Hyun Kim

## Abstract

Nicotinic acid adenine dinucleotide phosphate (NAADP) is an obligate driver of calcium signaling whose formation from other metabolites of nicotinamide adenine dinucleotide (NAD^+^) has remained elusive. *In vitro*, CD38-mediated NAADP synthesis requires an acidic pH and a nonphysiological concentration of nicotinic acid (NA). We discovered that the type II membrane form of CD38 catalyzes synthesis of NAADP by exchanging the nicotinamide moiety of nicotinamide adenine dinucleotide phosphate (NADP^+^) for the NA group of nicotinic acid adenine dinucleotide (NAAD) inside endolysosomes of interleukin 8 (IL8)-treated lymphokine-activated killer cells. Upon IL8 stimulation, cytosolic NADP^+^ is transported to acidified endolysosomes via connexin 43 via cAMP-EPAC-RAP1-PP2A signaling. Luminal CD38 then performs a base exchange reaction with the donor NA group deriving from NAAD, produced by newly described endolysosomal activities of NA phosphoribosyltransferase and NMN adenyltransferase 3. Thus, the membrane organization of endolysosomal CD38, a signal-mediated transport system for NADP^+^ and luminal NAD^+^ biosynthetic enzymes integrate signals from a chemokine and cAMP to specify the spatiotemporal mobilization of calcium to drive cell migration.

## INTRODUCTION

As helpful as the one gene-one enzyme hypothesis has been to definition of encoded biological functions, the multiplicity of enzymatic activities encoded by single genes and their insufficiency to confer key functions has frustrated molecular dissection of biological complexity. Calcium mobilization from acidic intracellular stores is mediated by formation of nicotinic acid adenine dinucleotide phosphate (NAADP) (1, 2), a metabolite related to nicotinamide adenine dinucleotide phosphate (NADP^+^) that can be formed *in vitro* by the membrane-associated CD38 protein performing a nucleobase exchange reaction between the nicotinamide (NAM) moiety of NADP^+^ and free nicotinic acid (NA) at acidic pH (3, 4). NAADP formation is stimulated by diverse stimuli including receptor binding and cell contact (5–7). In turn, high affinity release of calcium from acidic compartments mediates subsequent release of calcium from the endoplasmic reticulum that is required for mobility, differentiation, immune and contractile functions in plant and animal tissues (8). Because CD38 forms two additional nicotinamide adenine dinucleotide (NAD^+^)-related compounds, cyclic ADP-ribose and ADP-ribose, that mobilize calcium (9, 10), adopts at least two membrane orientations in multiple membrane compartments (11), and because NA is generally considered a bacterial NAD^+^ metabolite without appreciable availability in NAADP producing cells (12), it has not been clear what substrates CD38 uses to form NAADP and how substrates and CD38 are transiently colocalized to effect signal-dependent NAADP production.

In lymphokine-activated killer (LAK) cells, binding of interleukin 8 (IL8) to its G-protein coupled receptor, CXCR1, leads to activation of guanylyl cyclase and protein kinase G, which leads to internalization of CD38 and cADPR production (9, 10). In turn, cAMP is produced followed by formation of NAADP in lysosome-related acidic organelles, resulting in IL8-driven cell migration in a manner that depends on EPAC, protein kinase A and RAP1 (13).

Here, we used cell fractionation to identify and reconstitute signal-dependent, CD38-dependent and NADP^+^-dependent production of NAADP. Here we show that IL8 causes association of V-ATPase and a cAMP-EPAC-RAP1-PP2A-mediated activation of connexin 43 (Cx43) to transport NADP^+^ into the lumen of endolysosomes. The other CD38 substrate is here identified as nicotinic acid adenine dinucleotide (NAAD) produced by endolysosomal activities of NA phosphoribosyltransferase and NMN adenyltransferase 3. Compartmentalization of NAAD synthesis and signal-mediated acidification and transfer of NADP^+^ are thereby required to form NAADP and transmit signals for calcium mobilization.

## RESULTS AND DISCUSSION

As shown in Fig. 1A, in LAK cells, binding of IL8 to CXCR1 leads to activation of guanylyl cyclase and PKG, which leads to internalization of CD38 and cADPR production (9, 10). In turn, cADPR-mediated calcium release induces NAADP production in lysosome-related acidic organelles, resulting in IL8-driven cell migration in a manner that depends on Epac, protein kinase A and Rap1 (13). To identify the organelles that mediate production of NAADP, we treated LAK cells with IL8 and measured metabolite and protein levels in each subcellular fraction. Though CD38 was present in all fractions, only fraction 4 showed IL8-dependent production of NAADP. Fraction 4 was positive for late endosomal and lysosomal markers, Rab7 and LAMP1 (Fig. 1B).

**Figure 1.**
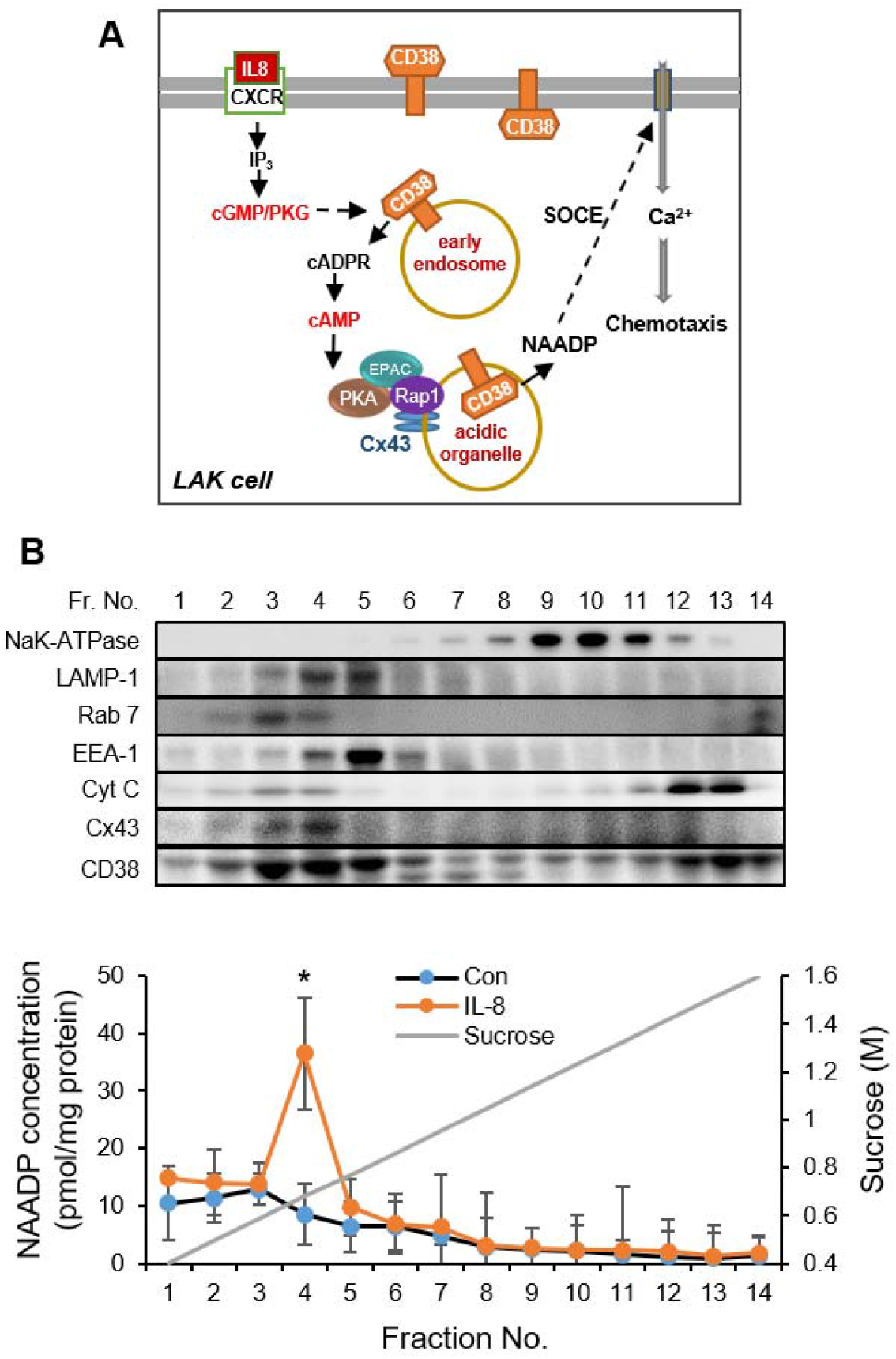
IL8 dependent NAADP formation occurs in a RAB7 and LAMP-1 rich subcellular compartment. (A) Schematic of IL8-driven NAADP production in LAK cells. (B) Subcellular fractionation of IL8-treated LAK cells identifies a unique CD38-containing fraction where NAADP is formed. Mean ± SEM of three independent experiments is shown. * *P* < 0.05, control versus IL8.

To determine the substrates for signal-dependent NAADP production, we quantified levels of NAD^+^ metabolites (14) in fractions before and after IL8 stimulation. Liquid-chromatography tandem mass spectrometry analysis revealed significantly decreased nicotinic acid adenine dinucleotide (NAAD) levels and increased NAD^+^ levels in fraction 4 after IL8 treatment (Fig. S1, A and B) while levels of nicotinic acid mononucleotide (NAMN), nicotinamide mononucleotide (NMN), and NADP^+^ (Fig. S1, C-E) remained unchanged. These findings suggest that NAAD might serve as the NA donor for formation of NAADP.

Because EPAC is an obligate mediator of NAADP formation in IL8-treated LAK cells (15), we tested whether NAADP formation increased when fraction 4 was treated with 8-pCPT-2′-O-Me-cAMP, a cAMP analog and selective EPAC activator. The activator increased NAADP formation in fraction 4 only when cytosol was provided (Fig. 2A), suggesting that import of an EPAC-dependent cytosolic component may be needed for NAADP synthesis. Given that NADP^+^ is an in vitro substrate for NAADP formation (3, 4), we tested whether NADP^+^ might substitute for cytosol in IL8-dependent NAADP formation by fraction 4. Moreover, because we identified high levels of Cx43, a known NAD^+^ transporter (16), in fraction 4 (Fig. 1B), we tested whether the EPAC-dependent cytosol requirement might be blocked by the Cx43 inhibitor, oleamide. As shown in Fig. 2A, NADP^+^ can substitute for cytosol in 8-pCPT-2′-O-Me-cAMP-stimulated NAADP formation in a manner that is sensitive to oleamide. In support of the Cx43 transport mechanism for EPAC-dependent NAADP formation, we showed that [^3^H]-NADP^+^ transport into fraction 4 is stimulated by the cAMP analog and blocked by the Cx43 inhibitor (Fig. 2B). Thus, IL8 and cAMP signaling drive a CD38 substrate into endolysosomes for NAADP formation.

**Figure 2.**
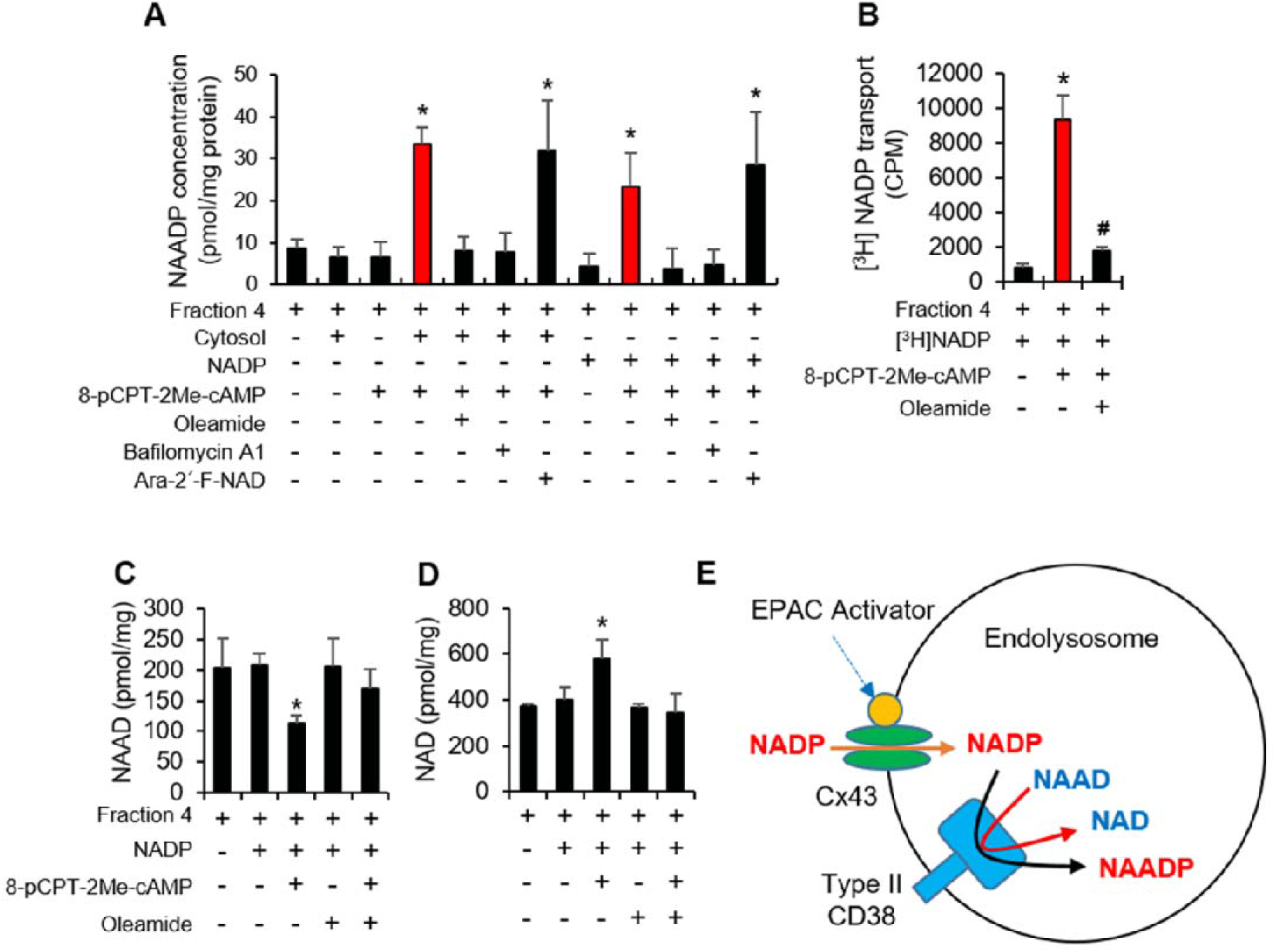
Signal-dependent NAADP formation depends on endolysosomal acidification, type II CD38 and EPAC-dependent import of NADP^+^. (A) Reconstituted NAADP formation from IL8-stimulated LAK cells occurs in a compartment that is acidified in a bafilomycin-inhibitable manner with cytosol that can be substituted by NADP^+^, which is transported in an EPAC- and Cx43-dependent fashion. NAADP synthesis was measured in fraction 4 after incubation with cytosol (60 μg) or NADP^+^ (100 μM) in presence or absence of EPAC activator, 8-pCPT-2Me-cAMP for 30 min at 37 °C. EPAC activator-induced NAADP synthesis was blocked by 100 μM oleamide (Cx43 inhibitor) and 1 μM bafilomycin A1 (lysosomal V-type H^+^-ATPase inhibitor) but not by 200 nM ara-2′-F-NAD^+^, a membrane-impermeable CD38 inhibitor. (B) NADP^+^ (100 μM) is transported (30 min, 37 °C). into Cx43-rich endolysosomes in a manner that depends upon EPAC stimulation (100 μM 8-pCPT-2Me-cAMP) and blocked by Cx43 inhibition (100 μM oleamide). (C and D) EPAC-stimulated NAADP formation is accompanied by stoichiometric depression of NAAD and formation of NAD^+^ quantified by LC-MS. (E) Schematic of NAADP formation in endolysosomes. Mean ± SEM of three independent experiments, *, effect of NADP^+^ and EPAC activator, and ^#^, effect of oleamide, *P* < 0.05.

The ability of CD38 to form NAADP rather than cADPR or ADPR depends on acidic pH (3, 4, 13). It has been established that endolysosomal acidification depends on assembly of the V_1_ and V_0_ subunits of V-ATPase (17). We discovered that fraction 4 contains V_0_ but not V_1_ subunits until cells are stimulated by IL8 (Fig. S2). Moreover, as shown in Fig. 2A, bafilomycin A1, an inhibitor of V-ATPase, completely blocked EPAC activator-induced NAADP synthesis. The requirement for organelle acidification and the involvement of Cx43-dependent NADP^+^ transport suggested that type II CD38, whose catalytic domain is in the lumen of organelles (11), must be responsible for NAADP production. We confirmed this by showing that ara-2′-F-NAD^+^, a membrane-impermeable CD38 inhibitor, is incapable of blocking EPAC activator-induced NAADP synthesis by fraction 4 (Fig. 2A).

Given that cellular NAAD levels were depressed by IL8 treatment (Fig. S1), we further analyzed NAD^+^ metabolite levels in fraction 4 in the presence of NADP^+^ before and after EPAC activator treatment. As in the reconstitution experiments with IL8-induced cell lysates (Fig. 1B), NAAD levels were significantly depressed and NAD^+^ levels were significantly increased after treating fraction 4 with the EPAC activator and NADP^+^ (Fig. 2C-D), with NAMN and NMN levels remaining unaltered (Fig. S3A-B). Changes in NAAD and NAD^+^ levels were blocked by oleamide (Fig. 2C-D). To exclude the possibility that NAAD was being converted to NAD^+^ by glutamine-dependent NAD^+^ synthetase (18), we reconstituted the proposed reaction (NAAD + NADP^+^ → NAADP + NAD^+^, Fig. 2E) with purified CD38 (Fig. S4). This reaction required only NAAD plus NADP^+^ under acidic conditions, produced stoichiometric amounts of NAADP and NAD^+^, and exhibited no requirement for glutamine or ATP.

A signal-dependent mechanism for NAADP formation must gate NADP^+^ import into an acidified CD38-containing organelle with high temporal specificity. Protein phosphatase PP2A is known to dephosphorylate Cx43 (19). Additionally, PP2A activity is positively modulated by cAMP in an EPAC and RAP1-dependent manner (20). Thus, we tested whether EPAC-RAP1-PP2A signaling might drive Cx43-mediated NADP^+^ transport. We discovered that IL8 stimulation increases association of EPAC, RAP1 and PP2A proteins with Cx43 (Fig. 3A) and CD38 (Fig. 3B). In whole cell extracts and fraction 4, both IL8 and EPAC activator increased the proportion of RAP1 in the GTP-bound form and its association with EPAC and PP2A (Fig. 3C). Moreover, we found that both IL8 and the EPAC activator increased PP2A enzymatic activity (Fig. S5A) and dephosphorylation of Cx43 at Ser-279 (Fig. S5B), suggesting that Cx43 dephosphorylation by PP2A induces NADP^+^ transport into endolysosomes. We further found that EPAC activator-induced NADP^+^ transport into fraction 4 organelles can be blocked by PP2A inhibitor okadaic acid (OA) and substituted by purified PP2A (Fig. 3D). These findings indicate that PP2A, once activated by EPAC and RAP1, dephosphorylates and gates Cx43 for NADP^+^, enabling production of NAADP by CD38 in endolysosomes upon IL8 treatment (Fig. 3E).

**Figure 3.**
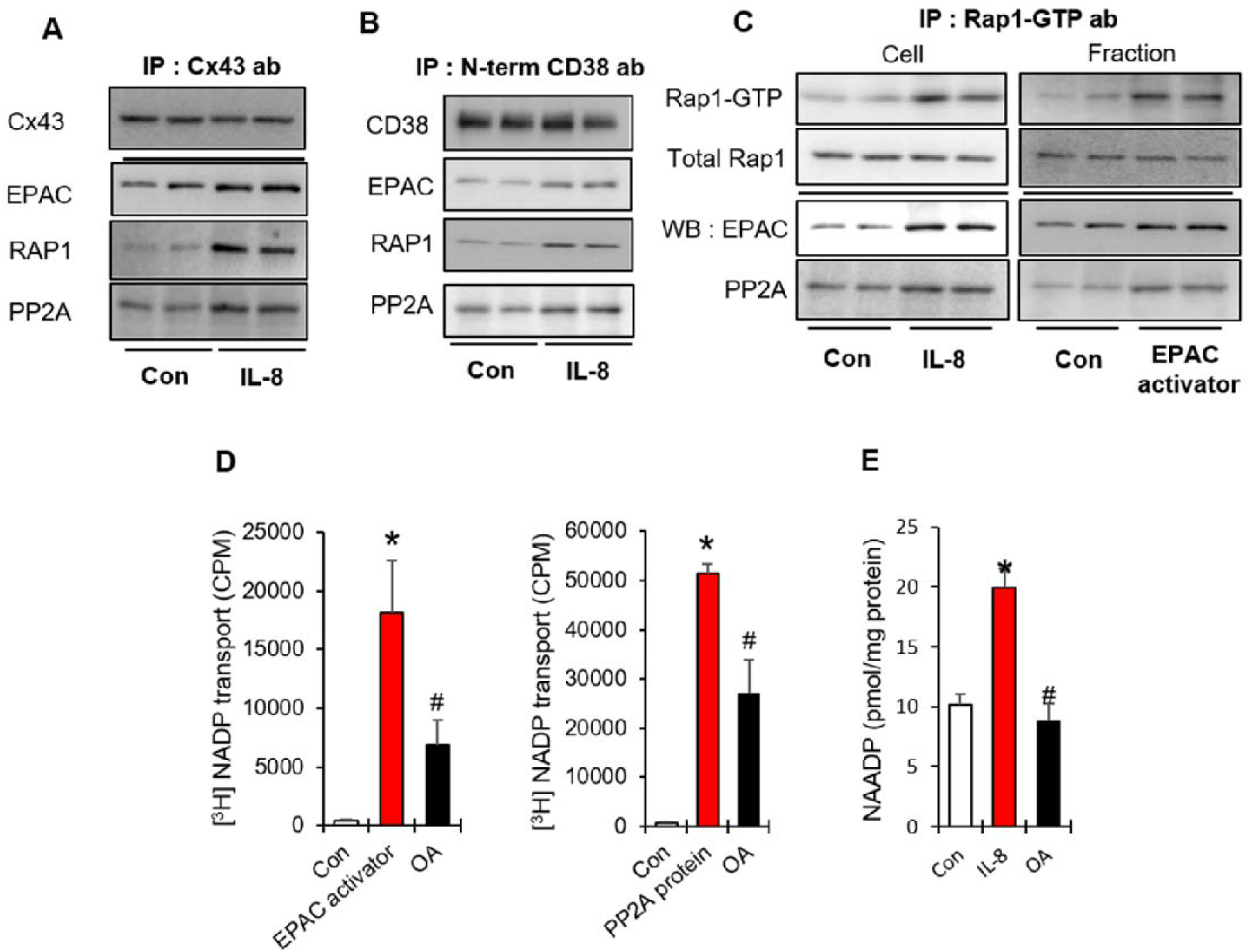
EPAC-RAP1-PP2A mediate NADP^+^ transport via Cx43 dephosphorylation. (A) IL8 induces association of EPAC-RAP1-PP2A complex with Cx43. (B) IL8 induces association of EPAC-RAP1-PP2A complex with CD38. (C) IL8 and EPAC activator increase association of RAP1-GTP with EPAC-PP2A in LAK cells and endolysosome fraction. (D) OA inhibits EPAC activator-induced NADP^+^ transport to endolysosome fraction. OA inhibits PP2A-induced NADP^+^ transport to the endolysosome fraction. (E) OA inhibits IL8-induced NAADP formation. Mean ± SEM of three independent experiments is shown. \**P* < 0.05, control versus EPAC activator; ^#^*P* < 0.05, EPAC activator versus OA.

Because NADP^+^ was identified as the only cytosolic NAD^+^ metabolite required to stimulate IL-8 or EPAC-signaled NAADP formation, and we identified NAAD as present in fraction 4 (Fig. S1A and Fig. 2C), we surmised that an NAMN adenylyltransferase activity must be localized in this fraction in order to produce the NAAD substrate. Three isoforms of NAMN/NMN adenyltransferase (NMNAT) are encoded by vertebrates (21). RT-PCR data showed that LAK cells express NMNAT1 and NMNAT3 (Fig. 4A). NMNAT1 and NMNAT3 have been termed the nuclear and mitochondrial isoforms respectively (21), though the null phenotype in mice is not consistent with this assignment for NMNAT3 (22). We used siRNAs to knock down NMNAT1 and NMNAT3 at the level of mRNA and protein (Fig. 4B). We discovered that IL8-induced NAADP formation in LAK cells was abolished by NMNAT3 knockdown but was unaffected by NMNAT1 knockdown (Fig. 4C). Similarly, EPAC activator-induced NAADP formation in fraction 4 depended on NMNAT3 but not NMNAT1 (Fig 4D). We further showed that fraction 4 contained high levels of NMNAT3 (Fig. S2). To further establish that NMNAT3 is responsible for NAAD synthesis in endolysosomes, we quantified levels of NAD^+^ metabolites in fraction 4 from LAK cells as a function of additions of EPAC activator and/or NADP^+^ and knockdown of NMNAT1 or NMNAT3. As expected, NAAD and NAD^+^ levels were significantly lower in samples from NMNAT3 knockdown cells when compared with those taken from NMNAT1 knockdown cells or control cells (Fig. 4E-F, Fig. S5). Consistent with known biosynthetic pathways (12), NAMN and NMN levels were significantly higher in the NMNAT3 and NMNAT1 knockdown LAK cells when compared to control cells (Fig. 4G-H, Fig. S6). Gallotannin has been characterized as an inhibitor of NMNAT with strong activity against the NMNAT3 isoform (21). We therefore examined the effect of gallotannin on EPAC activator-induced NAADP synthesis in fraction 4. Consistent with the hypothesis that an endolysosomal NMNAT3 activity is required to produce the NAAD substrate for NAADP synthesis, we found that gallotannin blocked EPAC activator-induced NAADP synthesis, resulting in significantly lower levels of NAAD and NAD^+^ and higher levels of NAMN and NMN in lysed fractions (Fig. S6).

**Fig. 4.**
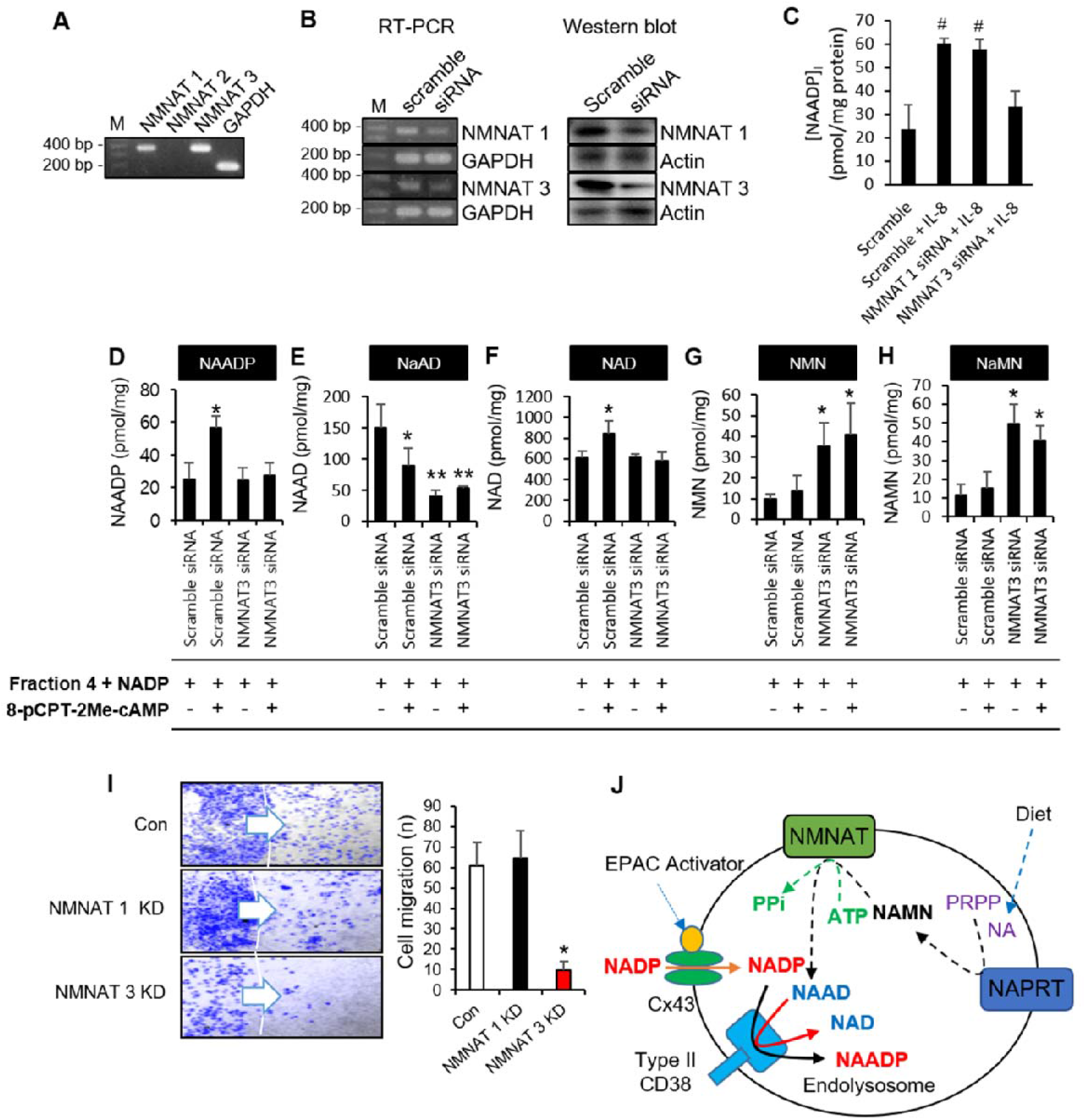
NMNAT3 is required to produce endolysosomal NAAD to effect IL8-driven NAADP-mediated chemotaxis. (A - C) mRNA expression of NMNAT1-3 in LAK cells as modulated by siRNA treatment and as assessed by Western blot. (D-H) IL8-induced NAADP formation in LAK cells depends on NMNAT3. NADP^+^ and EPAC activator-dependent nucleotide levels in fraction 4 depend on specific NMNAT isozymes. Endolysosomal concentrations of NAADP, NAAD, NAD^+^, NMN, and NAMN were determined by LC-MS after incubation (30 min at 37 °C) with 100 μM NADP^+^ plus or minus 8-pCPT-2Me-cAMP as a function of NMNAT3 status. The data indicate that NMNAT3 is required for NAAD formation and nonaccumulation of both NAMN and NAD^+^. Upon EPAC stimulation, CD38 converts NADP^+^ and NAAD to NAADP plus NAD^+^. (I) IL8-induced chemotaxis of LAK cells is sensitive to NMNAT3 KD. (J) Schematic of formation of NAAD via NAPRT and NMNAT3 in endolysosomes. Mean ± SEM of three independent experiments. ^#^*P* < 0.05 versus IL8 treated LAK cells; \**P* < 0.05, versus fraction 4 of scramble siRNA with 8-pCPT-2Me-cAMP; \*\**P* < 0.05 versus fraction 4 of NMNAT 3 siRNA.

Having established that the CD38 substrate NAAD is produced by endolysosomal NMNAT3, we asked the how the NMNAT substrate, NAMN, is synthesized. NAMN is synthesized either by *de novo* biosynthetic enzymes or by salvage of nicotinic acid (12), As shown in Fig. S7A, knockdown of the first and last *de novo* enzymes IDO and QPRT reduced IL8-induced NAMN abundance in fraction 4. In addition, the nicotinic acid salvage enzyme NAPRT (23) is abundant in fraction 4 (Fig. S7B) and NAAD synthesis was reconstituted from immunoprecipitated NAPRT from fraction 4 in a reaction with NA, PRPP, ATP and recombinant NMNAT (Fig. S7C).

Collectively these data establish that endolysosomes contain both the *de novo* and NA salvage enzymes for synthesis of one CD38 substrate, NAAD, the signal-dependent import machinery for the other substrate, NADP^+^, and a signal-dependent compartment acidification mechanism for NAADP formation from NADP^+^ and NAAD for type II CD38 activity. Earlier, it was shown that LAK cells from CD38 knockout mice cannot be induced to migrate by IL8 treatment (13). To test whether the proposed NAADP synthesis pathway is functionally important, we treated control, NMNAT1 and NMNAT3 knockdown cells with IL8 and assayed cell migration. As shown in Fig 4I, though both of these treatments depress total cellular NAAD and NAD^+^, only the NMNAT3 knockdown depressed cellular migration.

Intriguingly, EPAC, RAP1, Cx43 and V-ATPase have each been found to be involved in cell migration and cancer metastasis through unknown mechanisms (24–28). We conclude that endolysosomal NAADP synthesis by type II CD38 integrates cell surface inputs through these membrane-associated proteins to effect calcium-dependent cell migration.

## Materials and Methods

### Animals

C57BL/6 mice were purchased from Orient (Seongnam Korea). Mice were inbred and kept in animal facilities at Chonbuk National University Medical School under specific pathogen-free conditions. All studies conformed to the Guide for the Care and Use of Laboratory Animals published by the US National Institutes of Health (NIH Publication No. 85-23, revised 1996). The entire project was reviewed and approved by the Institutional Animal Care and Use Committee of the Chonbuk National University Medical School (CBNU 2018-069).

### Preparation of LAK cells

LAK cells were prepared as described previously (9, 10). Briefly, spleens of mice were minced harvested. RBCs were lysed by incubation with RBC lysis buffer (0.15 M NH_4_Cl, 10 mM KHCO_3_, 0.1 mM EDTA, pH 7.2) and the cells were thereafter washed with serum-free RPMI1640 twice. The RBC-removed cell preparations were incubated on a nylon-wool column at 37 °C for 1 h in a 5 % CO_2_ incubator to remove B lymphocytes and macrophages. Nylon-wool nonadherent cells were collected and incubated at 2 × 10^6^ cells/ml density in a culture media containing 3000 IU/ml IL2 in a 5 % CO_2_ incubator at 37°C. Culture media was RPMI 1640 supplemented with 10% FBS, 0.25 μg/ml amphotericin B, 10 U/ml penicillin G, 100 μg/ml streptomycin, 1 mM L-glutamine, 1 % nonessential amino acids, and 50 μM 2-mercaptoethanol. After incubation for 4 days, floating cells were removed, and adherent cells were cultured in the same media containing 3000 IU/ml IL2. LAK cells induced by IL2 for 8-10 days were used throughout the study.

### Subcellular fractionation of LAK cells and Western blotting

Organelles were separated as described (13). LAK cells were homogenized with a 30G syringe in 1 ml of a solution consisting of 0.25 M sucrose, 10 mM HEPES, pH 7.4 and 1 mM PMSF (homogenate buffer). The homogenate was then centrifuged at 1000 × *g* for 10 min at 4 °C and the supernatant was loaded on the top of a continuous 0.4-1.6 M sucrose gradient containing 1 mM EDTA, 10 mM HEPES, pH 7.4. After centrifugation at 100,000 × *g* for 20 h in a SW40Ti rotor (Beckman), 14 fractions were collected from the bottom of the tube. For Western blot analysis, an equal volume of 20 μl of each fraction were separated on either 10 % or 13 % SDS-PAGE and transferred to nitrocellulose membrane. The membrane was blocked in blocking buffer (10 mM Tris-HCl, pH 7.6, 150 mM NaCl, 0.05 % Tween 20) containing 5 % nonfat dry milk for 2 h at room temperature and then incubated with anti-NaK-ATPase (1:2500; Santa Cruz Biotechnology), anti-LAMP-1 (1:2500; BD Biosciences), anti-Rab-7 (1:2500; Santa Cruz Biotechnology), anti-EEA-1 (1:2500; Santa Cruz Biotechnology), anti-Cytochrome C (1:2500; BD Biosciences), anti-Cx43 (1:2500; Santa Cruz Biotechnology), anti-CD38 (1:2500; Santa Cruz Biotechnology), anti-V-ATPase v0 (1:2,500; Santa Cruz Biotechnology), anti-V-ATPase v1 (1:2500; Santa Cruz Biotechnology), or anti-NMNAT3 (1:2500; Santa Cruz Biotechnology) antibodies in the blocking buffer overnight at 4 °C. Blots were then washed with blocking buffer and incubated with HRP-conjugated anti-rabbit, anti-rat, anti-goat or anti-mouse antibodies (1:5000; Enzo Life Sciences) in the blocking buffer at room temperature for 2 h. The immunoreactive proteins with respective secondary Abs were determined using an enhanced chemiluminescence kit (Amersham Pharmacia Biotech) and exposed to an LAS 4000 Image Reader Lite (Fujifilm, Japan).

### Measurement of NAADP Concentration

Level of NAADP was measured using a cyclic enzymatic assay as described (29). Briefly, samples were treated with 0.2 ml of 0.6 M perchloric acid to stop the reaction. Precipitates were removed by centrifugation at 20,000 × *g* for 10 min. Perchloric acid was removed by mixing the aqueous sample with a solution containing three volumes of 2 M KHCO_3_. After centrifugation at 15,000 × g for 10 min, the aqueous layer was collected and neutralized with 20 mM sodium phosphate (pH 8). To remove all contaminating nucleotides, the samples were incubated with the following hydrolytic enzymes overnight at 37 °C: 2.5 units/ml apyrase, 0.125 unit/ml NADase, 2 mM MgCl_2_, 1 mM NaF, 0.1 mM PP_i_, 0.16 mg/ml NMN-AT in 20 mM sodium phosphate buffer (pH 8.0). Enzymes were removed by filtration using Centricon-3 filters. After the hydrolytic treatment, alkaline phosphate (10 units/ml) was added to convert NAADP to NAAD overnight at 37 °C. The alkaline phosphate was removed by filtration using Centricon-3 filters. The samples were further incubated with the cycling reagent containing 2% ethanol, 100 μg/ml alcohol dehydrogenase, 20 μM resazurin, 10 μg/ml diaphorase, 10 μM riboflavin 5’-phosphate, 10 mM nicotinamide, 0.1 mg/ml BSA, and 100 mM sodium phosphate (pH 8.0) at room temperature for 4 h. An increase in the resorufin fluorescence was measured at 544 nm excitation and 590 nm emission using a fluorescence plate reader (Molecular Devices Corp., Spectra-Max GEMINI). Known concentrations of NAADP were used to generate a standard curve.

### Measurement of [^3^H]NADP^+^ transport

[^3^H]NADP^+^ transport was performed as described (30) with modifications. Briefly, fraction 4 was incubated with 100 μM [^3^H]NADP and 100 μM 8-pCPT-2Me-cAMP for 30 min and samples were separated by a rapid oil-stop method (31). Transported [^3^H]NADP^+^ was quantified by scintillation counting (PerkinElmer Life Sciences).

### Measurement of NAAD, NAD^+^, NMN and NAMN

NAAD, NAD^+^, NMN, and NAMN levels were measured using LC-MS/MS as described (32). Briefly, organelle fraction samples were diluted with HEPES buffer (20 mM HEPES pH7.4, 10 mM KCl, and 2 mM MgCl_2_) that sucrose concentration is 0.25 M or less. Supernatant were removed by centrifugation at 15,000 xg for 10 min. Pellet were treated with 5 % trichloroacetic acid under sonication, and precipitates were removed by centrifugation at 20,000 × *g* for 10 min. Supernatants were loaded onto a Waters ACQUITY UPLC system coupled to a Waters Xevo TQ-S mass spectrometer and separated using a BEH Amide column (Waters ACQUITY UPLC BEH Amide, 130 Å, 1.7 μm, 2.1 mm × 50 mm). The column was equilibrated with 100 % buffer B (90 % acetonitrile/10 % 50 mM ammonium formate), and eluted with a 5-min gradient to 60 % buffer A (10 mM ammonium formate in water) at a flow rate of 0.5 ml/min. The following parameters were used for MS analysis: cone gas, 150 l/h; nebulizer, 7 Bar; and desolvation temperature, 350. The confirmation ion transitions for quantification were m/z 665→542 for NAAD, m/z 664→542.03 for NAD^+^, m/z 335.06→123.03 for NMN, and m/z 336→124 for NAMN.

### Measurement of NADP^+^

Level of NADP^+^ was determined by absorbance at 340 nm of extracts by the A_2_ - A_1_ subtractive method (33).

### Knockdown of NMNAT1, NMNAT3, IDO and QPRT by small interference RNA (siRNA)

NMNAT1, NMNAT3 and scrambled siRNA were purchased from Santa Cruz Biotechnology, QPRT siRNA was purchased from ThermoFisher, and the IDO siRNA referred was as described (34). LAK cells (2 × 10^5^ cells) were cultured in an antibiotic-free growth media supplemented with fetal bovine serum. After 24 h, LAK cells were transfected with 60 pmol of siRNA oligonucleotides specific to NMNAT1 and NMNAT3 using Lipofectamine RNAi/MAX (Invitrogen, Carlsbad, CA, USA) without antibiotics according to the instructions of the manufacturer. After 48 h of transfection, cells were prepared for examination.

### Reverse Transcriptase PCR (RT-PCR)

Total RNA was isolated from LAK cells using the Hybrid-R^™^ Kit (GeneAll). cDNA was synthesized by reverse transcription from 1 μg total RNA using a ImProm-II^™^ Reverse Transcription System (Promega). Primers for NMNAT1, NMNAT2, NMNAT3 and GAPDH were 5’-TGCCCAACTTGTGGAAGATG-3’ (NMNAT1 forward primer), 5’-AATGGTTGTGCTTGGCCTCT-3’ (NMNAT1 reverse primer), 5’-TGGAGCGCTTCACTTTTGTAG-3’ (NMNAT2 forward primer), 5’-GATGTACAGCTGACTCTTGA-3’ (NMNAT2 reverse primer), 5’-AGCACTGCCAGAGTTGAAAC-3’ (NMNAT3 forward primer), 5’-CTTTCCAGGAACCGTCATTG-3’ (NMNAT3 reverse primer), 5’-CGTGGAGTCTACTGGTGTCTT-3’ (GAPDH forward primer) and 5’-GTTGGTGGTGCAGGATGCATT-3’ (GAPDH reverse primer). Thirty cycles of amplification were carried out as follows: 30 sec at 94 °C; 30 sec at 62 °C; 30 sec at 72 °C, followed by 5 min elongation at 72 °C.

### NAADP transport

NAADP transport experiments were performed as described (30) with minor modifications. Briefly, Fraction 4 was incubated with inhibitors and 100 μM 8-pCPT-2Me-cAMP for 30 min at 37 °C and the reaction mixes were separated by rapid oil-stop (31). NAADP was measured by a cyclic enzymatic assay.

### Immunoprecipitation

Organelle fraction 4 was incubated 100 μM 8-pCTP-2Me-cAMP for 30 min at 37 °C and then lysed with an ice-cold lysis buffer (20 mM Tris-HCl, pH 7.2, 2 % NP-40, 150 mM NaCl, 1 mM phenylmethylsulfonylfluoride, 10 μg/ml leupeptin, 10 μg/ml pepstatin, and 10 μg/ml aprotinin). After centrifugation at 20,000 × *g* for 10 min, supernatants were taken. For immunoprecipitation, lysate precleared with protein G-agarose was incubated with anti-CD38 mAb overnight at 4 °C, and then further incubated with protein G-agarose at 4 °C for 1 h. The immunoprecipitates were washed four times with lysis buffer and then used for western blot and HPLC.

### Cell migration assay by agarose spot assay

Chemokine-induced cell migration assay was performed as described (35). 0.1 g of low-melting point agarose (Invitrogen) was placed into a 100 mL beaker and diluted into 20 mL PBS to make a 0.5% agarose solution. This was heated on a hot plate in the cell culture hood until boiling, swirled to facilitate complete dissolution, and then taken off of the heat. When the temperature cooled to 40°C, 90 μL of agarose solution was pipetted into a 1.5-mL Eppendorf tube containing 10 μL of PBS with or without IL8 (Sigma). Ten-microliter spots of agarose containing IL8 were pipetted, using cut pipet tips (Cat. no. 70.760.211; Sarstedt, Leicester, UK), as rapidly as possible onto 35-mm glass-bottomed dishes (SPL, Korea), and allowed to cool for ~5 min at 4°C. Four spots per dish were pipetted, two containing IL8 and two containing only PBS. At this point cells were plated into spot-containing dishes in the presence of 10% FBS cell culture media and allowed to adhere for 4 h. Cells were transferred into cell culture media or Hanks’ balanced salt solution (Invitrogen) with 10 mM HEPES (Sigma-Aldrich, Dorset, UK) for time-lapse studies with 0.1% FBS, replaced into the incubator for 4 h, and analyzed by microscopy. Imaging was performed on a Nikon TE300 inverted microscope with a 10x objective (Nikon, Surrey, UK), and for each spot we recorded the field that contained the highest apparent number of motile cells penetrating furthest underneath the agarose spot.

### HPLC

NAADP was quantified as described (36). Nucleotide synthesis reactions from NADP^+^ and NAAD substrates contained 20 mM sodium acetate (pH 4.5), 1 mM calcium chloride, 1 mM magnesium chloride. For HPLC analysis, nucleotides were separated on a column (3 × 150 mm), packed with AGMP-1 resin (Bio-Rad Laboratories). Samples (100 μl) were injected onto a column equilibrated with water. The bound material was eluted with a concave-up gradient of trifluoroacetic acid, which increased linearly to 2% at 1.5 min and to 4, 8, 16, 32 and 100% (150 mM trifluoroacetic acid) at 3, 4.5, 6.0, 7.5 and 7.51 min respectively. The flow rate was 4 ml/min. Nucleotides were detected by an absorbance at 254 nm. Detection of NAMN, NAD^+^, NAAD was performed by reverse phase HPLC method as described (37). Nucleotides were separated on a BDS hyperseal C18 column (Thermo). Samples (100 μl) were injected onto a column equilibrated with 0.05 M phosphates buffer. The HPLC is run at a flow rate of 1 mL/min with 100% buffer A (0.05 M Phosphate Buffer) from 0-5 min, a linear gradient to 95% buffer A/5% buffer B (100% methanol) from 5-6 min, 95% buffer A/5% buffer B from 6-11 min, a linear gradient to 85% buffer A/15% buffer B from 11-13 min, 85% buffer A/15% buffer B from 13-23 min, a linear gradient to 100% buffer A from 23-24 min, and 100% buffer A from 24-30 min. The flow rate was 1 ml/min. Nucleotides were detected by absorbance at 261 nM.

### PP2A activity

PP2A activity were measured by PP2A phosphatase activity assay kit from Merck Millipore (PP2A immunoprecipitation phosphatase assay kit, 17-313). IL8 and 100 μM 8-pCTP-2Me-cAMP or 1 μM OA-treated LAK cells or organelle fraction 4 was lysed with extraction buffer. PP2A activity was measured using an ELISA reader (Bio-rad xMARK microplate spectrometer) at a wavelength of 650 nm.

## QUANTIFICATION AND STATISTICAL ANALYSIS

Data are expressed as means ± SEM. Statistical comparisons were performed with analysis of variance (ANOVA) using Sigma plot. Significant differences between groups were determined with the unpaired Student’s t test. Statistical significance was set at *P* < 0.05

## AUTHOR CONTRIBUTIONS

U-HK designed the experiments. TSN and DRP performed experiments with help from SYR and TGW. Data were analyzed by U-HK, HTC, TSN, DRP and CB. The manuscript was written by U-HK and CB with help from TSN, DRP and HTC.

## ACKNOWLEDGMENTS

We thank Chansu Park for critical reading of the manuscript. Funding: This work was supported by Korean National Research Foundation Grant 2012R1A3A2026453 to U-HK, 2014R1A6A1030318 to HTC, and funds from the Roy J. Carver Trust to CB.

