## Supplemental Figures for "IL8 Drives an Elaborate Signal Transduction Process for CD38 to Produce NAADP from NAAD and NADP^+^ in Endolysosomes to Effect Cell Migration"

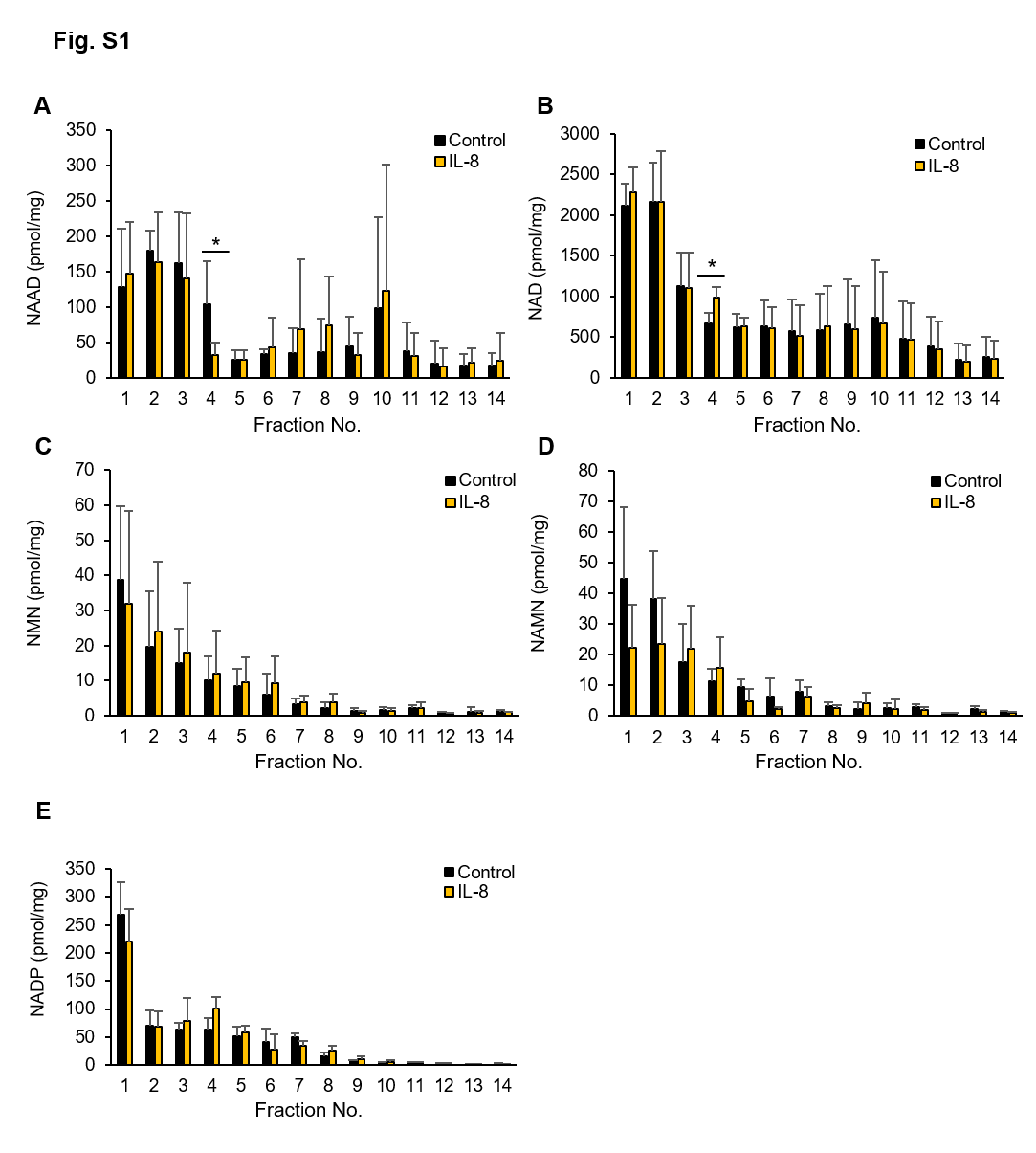

**Fig. S1.** Changes of (A) NAAD, (B) NAD^+^, (C) NMN, (D) NAMN and (E) NADP^+^ levels in subcellular fractions of LAK cells before and after treatment with IL8. Levels of NAAD, NAD^+^, NMN, NAMN and NADP^+^ were measured by LC-MS/MS method. LAK cells were stimulated with or without 10 pM IL8 for 10 sec and cell lysates were fractionated by sucrose gradient centrifugation. * p < 0.05, versus control fraction. Mean ± SEM of three independent experiments is shown.

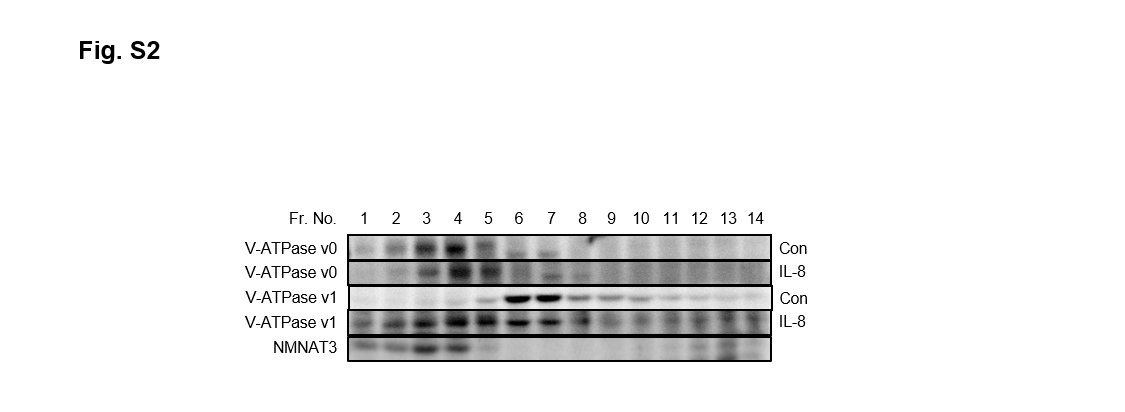

**Fig. S2.** Subcellular fractions were analyzed for V-ATPase V_0_ and V-ATPase V_1_ by Western blotting. LAK cells were stimulated with or without 10 pM IL8 for 10 sec and cell lysates were fractionated by sucrose gradient centrifugation.

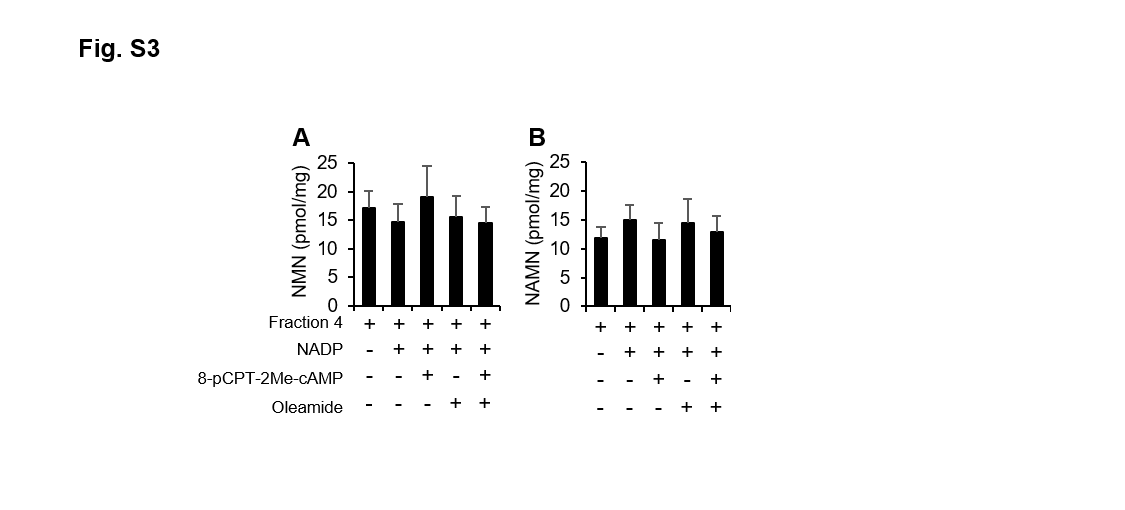

**Fig. S3.** Inhibition of Cx43 has no effect on NMN and NAMN levels in endolysosome. Fraction 4 was stimulated by 100 µM 8-pCPT-2Me-cAMP for 30 min at 37℃ in presence or absence of Cx43 inhibitor, oleamide. Levels of NMN and NAMN were measured by LC-MS/MS method.

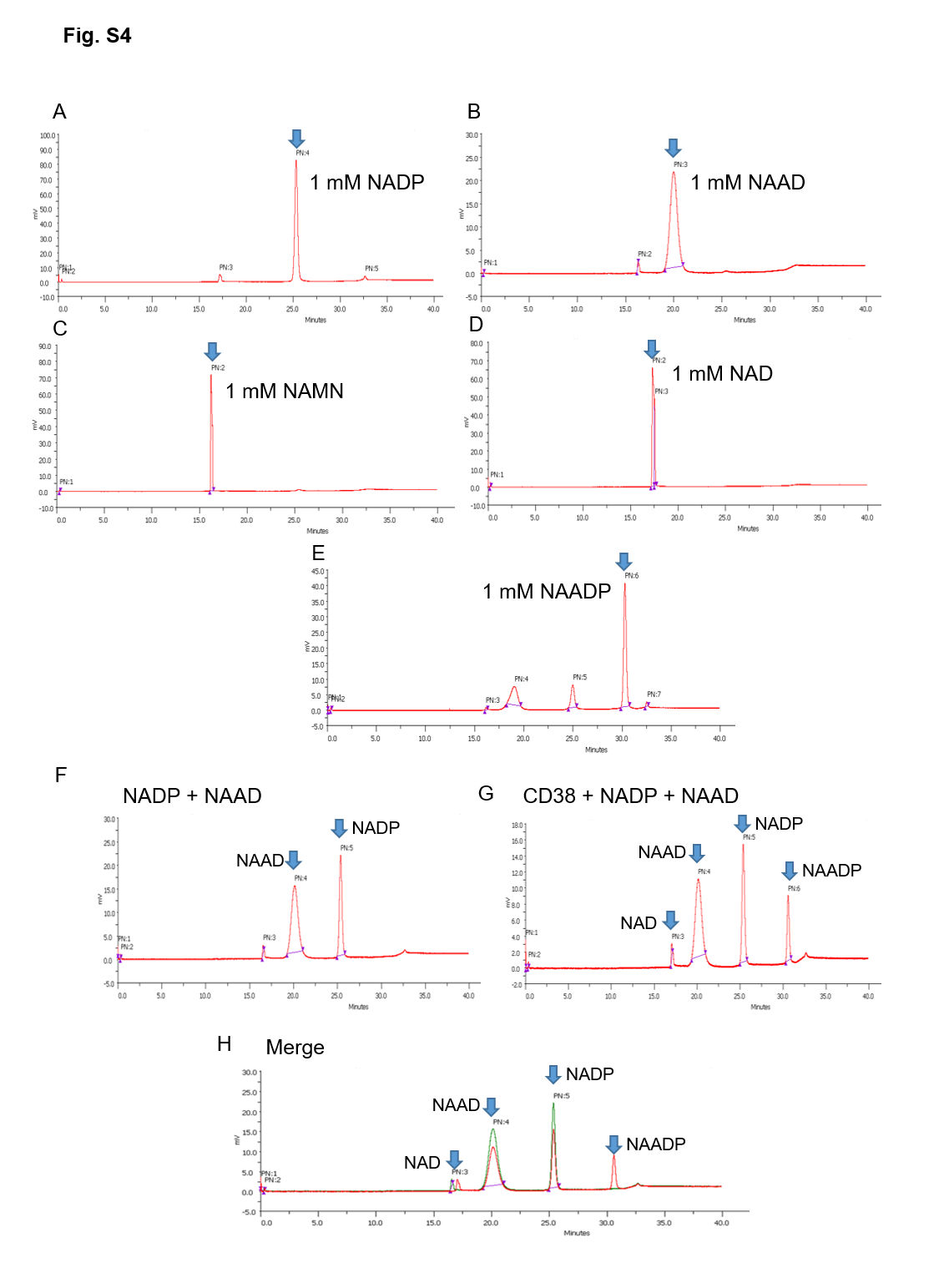

**Fig. S4.** NAADP is produced from NADP^+^ and NAAD through base exchange reaction. (A-E) Retention times were compared with known standards (1 μM). (F-H) HPLC analysis was performed following treatment of NADP^+^ (1 μM), NAAD (1 μM) with purified CD38 at acidic condition (pH 4.5).

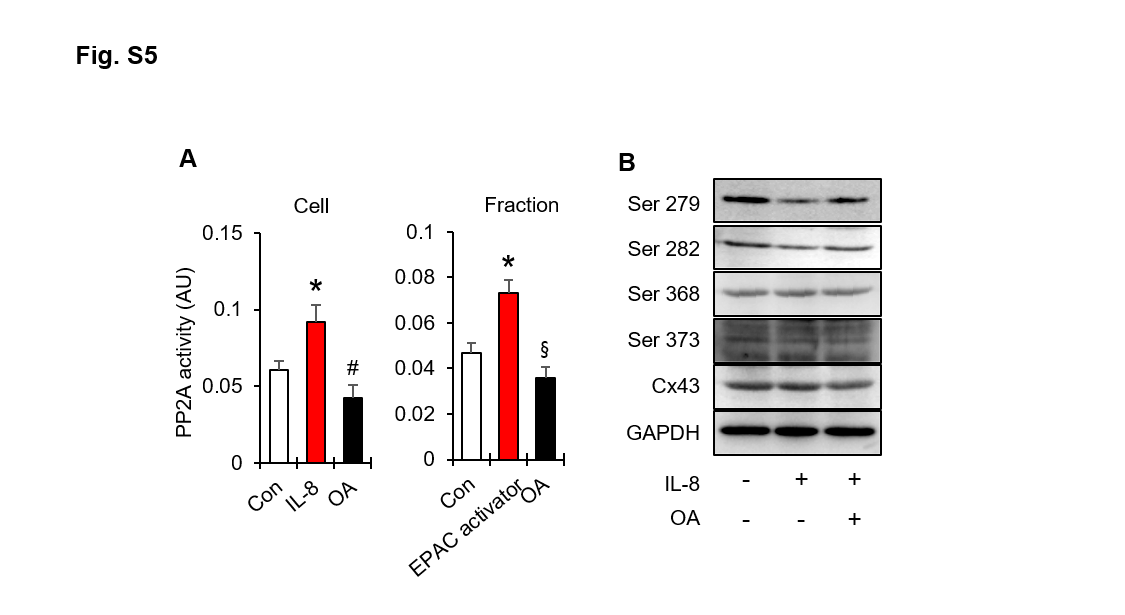

**Fig. S5.** Treatment of IL8 or EPAC activator increases Cx43 dephosphorylation in LAK cell or endolysosomal fraction. (A) 10 pM of IL8 and 100 µM EPAC activator increase PP2A activity in LAK cells or endolysosome. Okadaic acid (OA) inhibits IL8- or EPAC activator-induced PP2A activation (B) IL8 increases Cx43 dephosphorylation at Ser279 and Ser282 via PP2A in LAK cell. * p < 0.05, versus control. ^#^ p < 0.05, versus IL-8 or EPAC activator treated conditions.

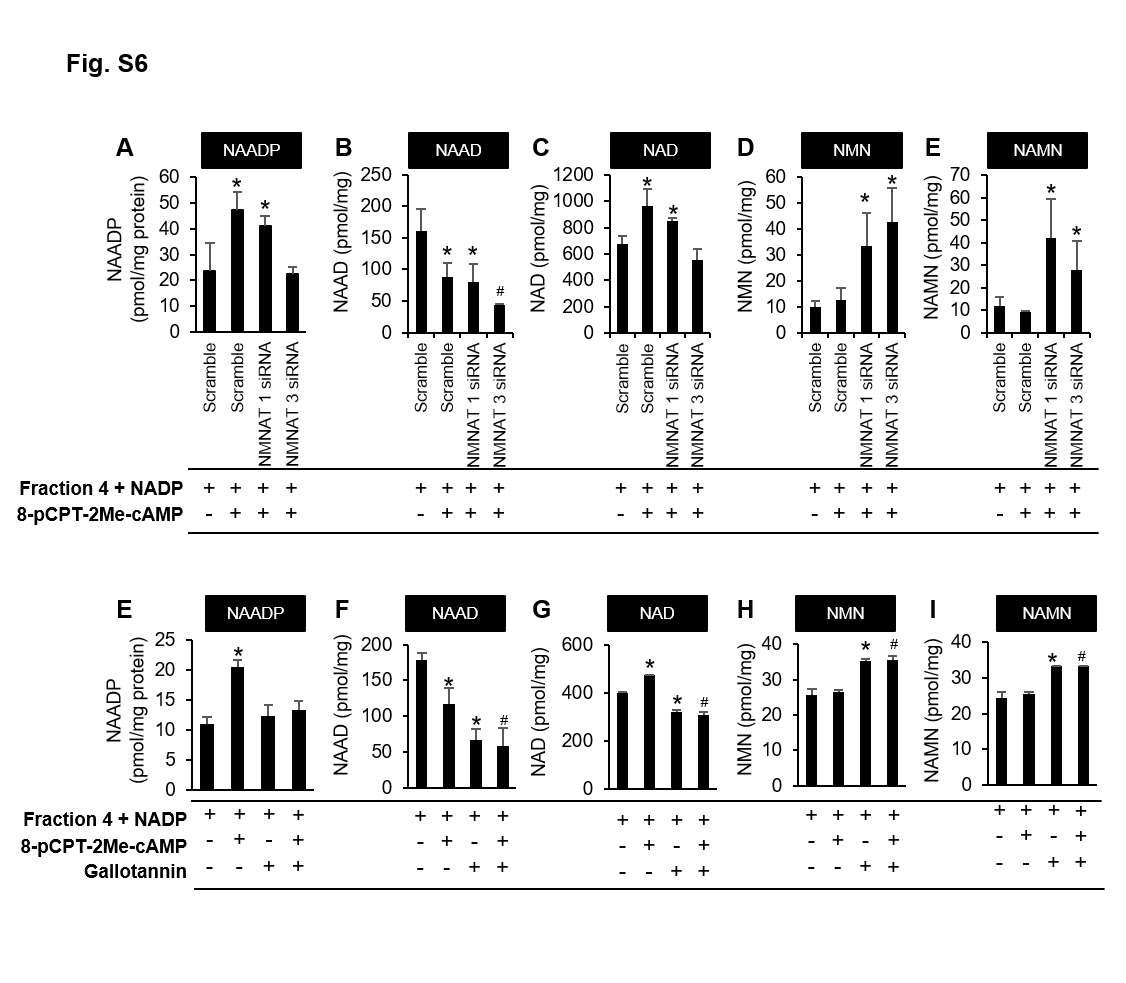

**Fig. S6.** Changes of NAADP, NAAD, NAD^+^, NMN, NAMN levels in endolysosome fraction of LAK cells before (A-E) and after treatment with gallotannin (E-I). Levels of NAAD, NAD^+^, NMN, and NAMN were measured by LC-MS/MS method. Fraction 4 was obtained by sucrose density centrifuge method from LAK cells incubated with or without 100 µM gallotannin (NMNAT inhibitor) for 30 min at 37℃. The effect of gallotannin on NAADP, NAAD, NAD^+^, NMN, and NAMN levels changes was compared with in the presence or absence of 100 µM NADP^+^ and 100 µM 8-pCPT-2Me-cAMP for 30 min at 37℃. * p < 0.05, versus fraction 4 with NADP^+^. ^#^ p < 0.05, versus fraction 4 with NADP^+^ and 8-pCPT-2Me-cAMP. Mean ± SEM of three independent experiments is shown.

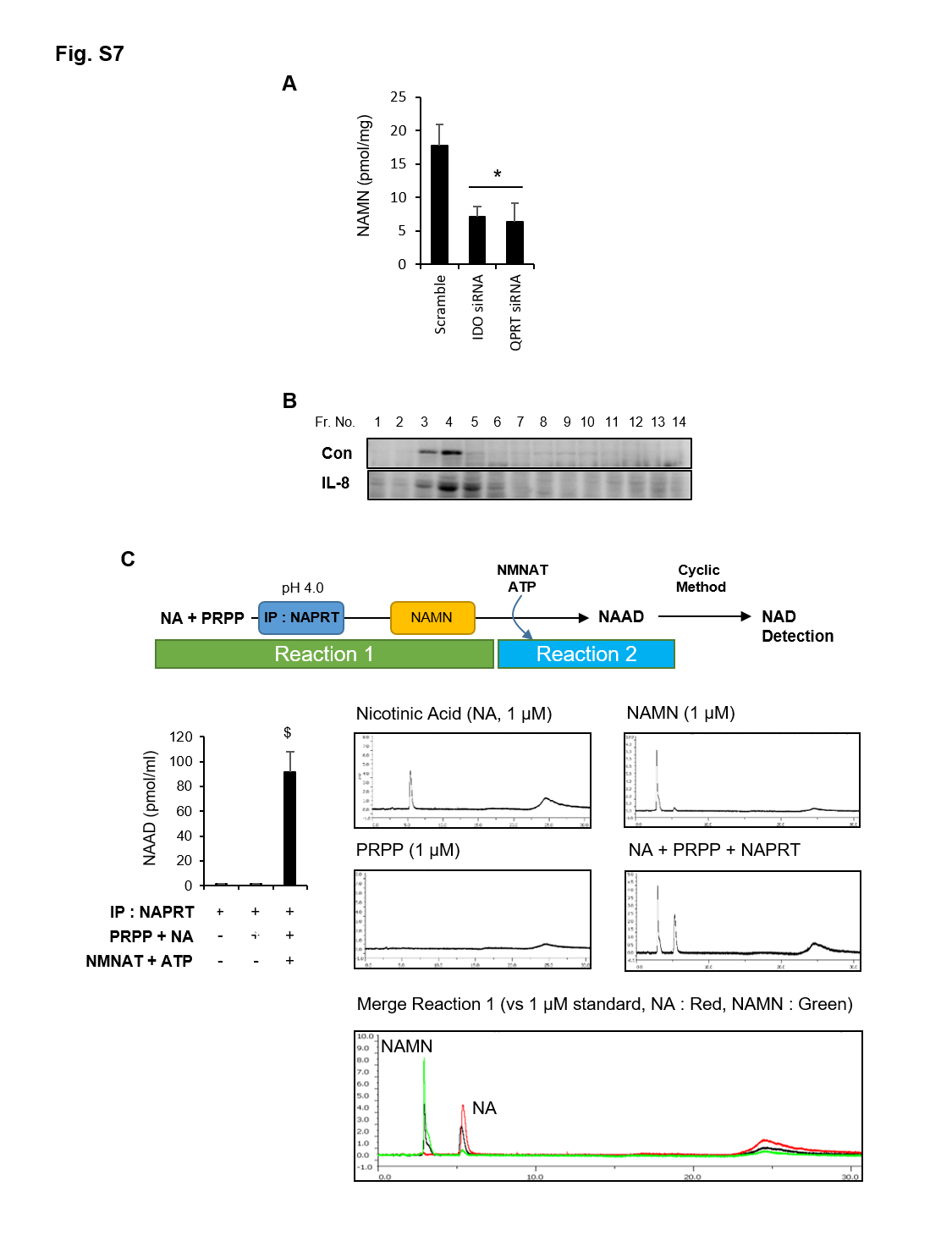

**Fig. S7.** NAMN synthesized by NAPRT in endolysosome. (A) NAMN level decreased by knock down of tryptophan metabolism regulator, indoleamine 2,3-dioxygenase (IDO) and quinolinic acid phosphoribosyltransferase (QPRT). * p < 0.05, versus scramble. (B) Expression of nicotinic acid phosphoribosyltransferase (NAPRT) in endolysosomal fraction. Control or IL8-treated LAK cell lysates were fractionated by sucrose gradient centrifugation. NAPRT was detected by Western blotting. (C) NAMN is synthesized with nicotinic acid (NA) and phosphoribosyl pyrophosphate (PRPP) via NAPRT, thereby being substrate for NAAD formation by NMNAT in endolysosome. NAPRT was immunoprecipitated from fraction 4 and incubated with 1 µM NA and 1 µM PRPP in acidic condition (pH 4.0). Resulting product, NAMN, is incubated with recombinant NMNAT and 1 µM ATP in same condition. All nucleotides were detected by HPLC method. ^$^ p < 0.05, versus without reactions.
